# “Characterization of spatiotemporal dynamics of binary and graded tonic pain in humans using intracranial recordings”

**DOI:** 10.1101/2023.03.08.531576

**Authors:** Rose M. Caston, Elliot H. Smith, Tyler S. Davis, Hargunbir Singh, Shervin Rahimpour, John D. Rolston

**Affiliations:** Department of Biomedical Engineering, University of Utah, 84112; Department of Neurosurgery, University of Utah, 84112; Interdepartmental Program in Neuroscience, University of Utah, 84112; Department of Neurosurgery, Brigham & Women’s Hospital, Harvard Medical School, 02115

## Abstract

Pain is a complex experience involving sensory, emotional, and cognitive aspects, and multiple networks manage its processing in the brain. Examining how pain transforms into a behavioral response can shed light on the networks’ relationships and facilitate interventions to treat chronic pain. However, studies using high spatial and temporal resolution methods to investigate the neural encoding of pain and its psychophysical correlates have been limited.

We recorded from intracranial stereo-EEG (sEEG) electrodes implanted in sixteen different brain regions of twenty patients who underwent psychophysical pain testing consisting of a tonic thermal stimulus to the hand. Broadband high-frequency local field potential amplitude (HFA; 70-150 Hz) was isolated to investigate the relationship between the ongoing neural activity and the resulting psychophysical pain evaluations. Two different generalized linear mixed-effects models (GLME) were employed to assess the neural representations underlying binary and graded pain psychophysics. The first model examined the relationship between HFA and whether the patient responded “yes” or “no” to whether the trial was painful. The second model investigated the relationship between HFA and how painful the stimulus was rated on a visual analog scale.

GLMEs revealed that HFA in the inferior temporal gyrus (ITG), superior frontal gyrus (SFG), and superior temporal gyrus (STG) predicted painful responses at stimulus onset. An increase in HFA in the orbitofrontal cortex (OFC), SFG, and striatum predicted pain responses at stimulus offset. Numerous regions including the anterior cingulate cortex, hippocampus, IFG, MTG, OFC, and striatum, predicted the pain rating at stimulus onset. However, only the amygdala and fusiform gyrus predicted increased pain ratings at stimulus offset.

We characterized the spatiotemporal representations of binary and graded painful responses during tonic pain stimuli. Our study provides evidence from intracranial recordings that the neural encoding of psychophysical pain changes over time during a tonic thermal stimulus, with different brain regions being predictive of pain at the beginning and end of the stimulus.

**Significance Statement:** We investigated the neural encoding of pain psychophysics across 16 brain regions during a continuous thermal stimulus in humans. Mixed-effects models were used to analyze trends across 20 human subjects. Using intracranial electrodes, we show a parametric relationship between behavioral responses and HFA during ongoing pain. We found that HFA in cognitive and emotional pain processing regions was closely associated with pain evaluation at the stimulus onset, end, or both. The neural encoding of subjective pain intensity, measured by a visual analog scale, differed from that of binary pain intensity. Perception and psychophysical correlates to pain depend on how patients are asked to evaluate it. Our findings provide evidence that HFA can serve as a neural marker within specific brain regions of behavioral pain responses, as measured by sEEG.

## Introduction

Pain is a complex and multi-faceted experience that encompasses sensory, emotional, and cognitive aspects (Tan and Kuner, 2021). The ability to feel pain is a crucial defense mechanism managed by several processing networks. Unlike other sensory modalities, pain does not appear to have a single primary brain region responsible for initial processing, as shown by multiple studies (Kucyi and Davis, 2015; Bastuji et al., 2016; Seymour, 2019; Caston et al., 2020). When pain persists without an apparent cause, it becomes a chronic condition that can devastate one’s quality of life. When pain arises without a noxious stimulus, it suggests a disruption in the connection between the stimulus and psychophysical correlates. Understanding this interaction could offer valuable information for managing chronic pain.

Much of what is known about pain processing has resulted from studies that induce pain in healthy volunteers using brief, millisecond-long stimuli with infrared lasers (Markman et al., 2013; Kim et al., 2015; Liu et al., 2015). Such work has revealed pain networks comprise of regions such as the somatosensory, insular, and prefrontal cortices (Davis et al., 2002; Wager et al., 2013; Asad et al., 2016; Xu et al., 2020; Mancini et al., 2022). Noninvasive imaging studies inducing tonic pain with thermodes suggest that ongoing pain activates similar brain regions as brief experimental stimuli (Schreckenberger et al., 2005; Owen et al., 2010; Wasan et al., 2011). However, it’s thought that experimental tonic pain also involves the medial prefrontal cortex and related projections, such as the anterior cingulate cortex (Marusak et al., 2016), nucleus accumbens (Baliki et al., 2010), hippocampus, periaqueductal gray, globus pallidus, and subthalamic nucleus (Etkin et al., 2011; Schulz et al., 2015; Ong et al., 2019). The involvement of these regions during experimental tonic pain may indicate a shift from sensory to emotional and cognitive encoding-based processes (Tan and Kuner, 2021). The spatiotemporal dynamics of these emotional and cognitive encoding-based processes are crucial in understanding how pain is shaped.

We sought to understand the psychological aspects of painful experiences with high spatiotemporal resolution recordings of neuronal population activity in human participants and statistical modeling of two types of psychophysical ratings. We hypothesized that regions involved in somatosensation and perception would have increased gamma activity if the subject later evaluated that stimulus as painful. At the end of the stimulus, we hypothesized that high gamma activity would dissipate from regions involved at the onset of the trial and instead would develop in regions involved in cognitive-emotional processing. Our results revealed that the inferior, middle and superior temporal gyrus (ITG, MTG, STG) predicted both binary and numerical psychophysical correlates of pain. Other regions, such as the cingulate cortex, orbitofrontal, hippocampus, and amygdala, had HFA amplitudes associated with higher numerical ratings of pain. We describe the utility of understanding how these regions interact and propose future studies of interest.

## Methods and Materials

### Patient Selection

Patients undergoing routine intracranial monitoring with sEEG for localization of epileptic foci were screened for inclusion in the study between 2020 and 2022. A total of 20 patients (12 female, 8 male) met the inclusion criteria (see below) and were included in the final data analysis. The sample size was determined from an a priori power analysis with an assumed effect size (f = 0.625), type I error probability (α= 0.05), and power threshold (0.80). The age range of the patient population was 21-66 years (mean 38 ± 10 years), and all patients had a diagnosis of drug-resistant epilepsy as determined by a multidisciplinary conference at our institution. All patients underwent sEEG implantation for clinical purposes. Participation in the research study was obtained through informed consent under a protocol approved by the University of Utah Institutional Review Board. To be eligible for the study, participants must have been 18 years of age or older, able to give informed consent, without any nerve damage in their arms or hands, able to communicate during the task, and not have any serious medical conditions such as bleeding disorders or cancer. Patients were excluded if their seizures interfered with data collection.

### Psychophysical Pain Task

Our research group recently developed and fabricated a thermoelectric device compatible with intracranial electrodes. The device and integrated software have been validated as a psychophysical pain task in healthy human subjects (Caston et al., 2023). The task consists of four main events (**Figure 1A**), which occur in a single trial: 1) The participant places their hand on the device for 10 seconds, 2) the participant removes their hand from the device, 3) the participant responds to whether the trial was perceived as painful, and 4) the participant rates the perceived pain and heat intensity on a scale of 0-10. Each of these events is time-locked to the ongoing intracranial recording. The start and end of each trial are time-locked using a capacitive touch sensor under the device’s surface. The patient used their dominant hand unless an intravenous access site was on that hand.

**Figure 1.**
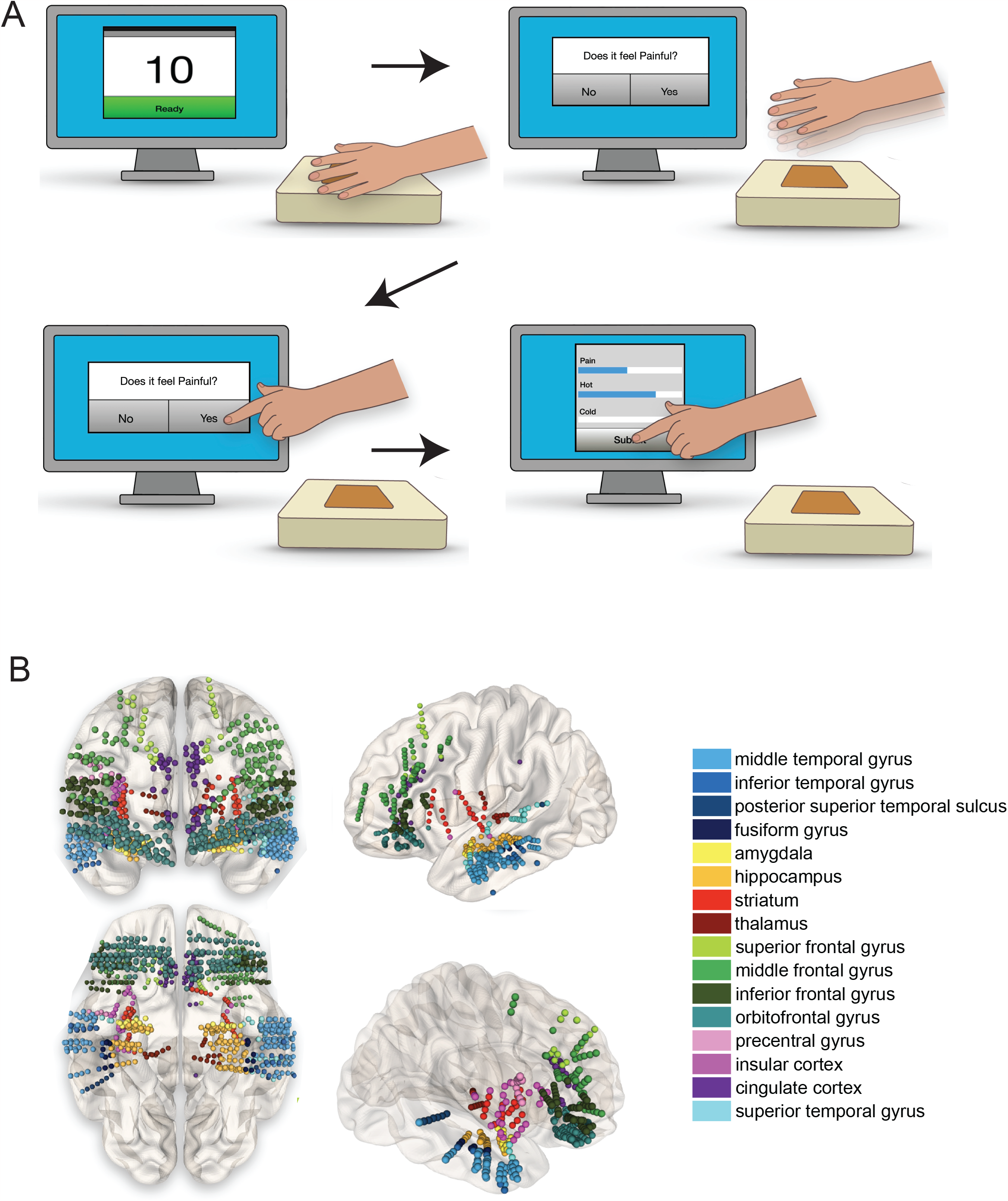
Psychophysical task paradigm and intracranial recording locations. 1A: Pain stimuli of varying intensity were applied to the hand’s palmar surface in patients with intracranial electrodes. The intensity of the stimulus was adjusted in each trial. One trial constituted having the hand on the stimulus for 10 seconds (top left). The patient removed their hand from the device (top right). After removing their hand, the patient responded to the question, “Was that painful?” on the touchscreen monitor (bottom left). After answering “yes” or “no,” the patient filled out how painful, hot, or cold the stimulus felt on a scale of 0-10 (bottom right). After submitting their answers, the stimulus was adjusted, and the sequence was repeated. 1B: A wide array of neural structures were sampled. Electrode locations for all 20 patients are shown according to the coordinates in MNI space. We analyzed electrodes within a known gray matter location of the Brainnetome Atlas. The electrode location also had at least three unique patients representing it within the GLME models. Electrode locations were colored according to the region of interest. While this figure shows electrode locations as right and left, we did not distinguish lateralization within the analysis.

The thermoelectric device also includes built in software that implements a psychophysical algorithm to estimate pain thresholds (QUEST Psychtoolbox, (Watson, 2017)). The algorithm incorporates the participant’s responses after each trial. This enables adaptation to the most likely pain threshold temperature. Participants completed at least 20 trials to allow adequate sampling for signal processing. The full description of the psychophysical pain task and its capability is reported elsewhere (Caston et al., 2023).

### Electrode Localization

The study utilized electrodes designed for sEEG. The locations of the electrodes were identified using each patient’s structural magnetic resonance imaging coregistered with postoperative high-resolution computed tomography using the LeGUI software package (https://github.com/Rolston-Lab/LeGUI) (Davis et al., 2021). The Brainnetome atlas was used within LeGUI to label the anatomical region based on the electrode locations (http://atlas.brainnetome.org) (Fan et al., 2016). The electrode labels were then converted to regional gyri by only keeping the first part of the alphanumerical atlas label. The distinction between the right and left hemispheres was also removed. For example, the Brainnetome label “Amyg_L_2_1” was collapsed to “Amyg.” Electrode locations were not included if they were within white matter or if at least three unique patients did not have electrodes in the same region.

Sixteen total areas were represented by 3 to 20 unique patients (**Figure 1B**). The number of patients and the average number of electrodes representing each brain area is shown in **Table 1**. The electrodes sampling the 16 brain regions were visualized in Montreal Neurological Institute (MNI) space using NeuroMArVL (https://github.com/mangstad/neuromarvl.

**Table 1.**
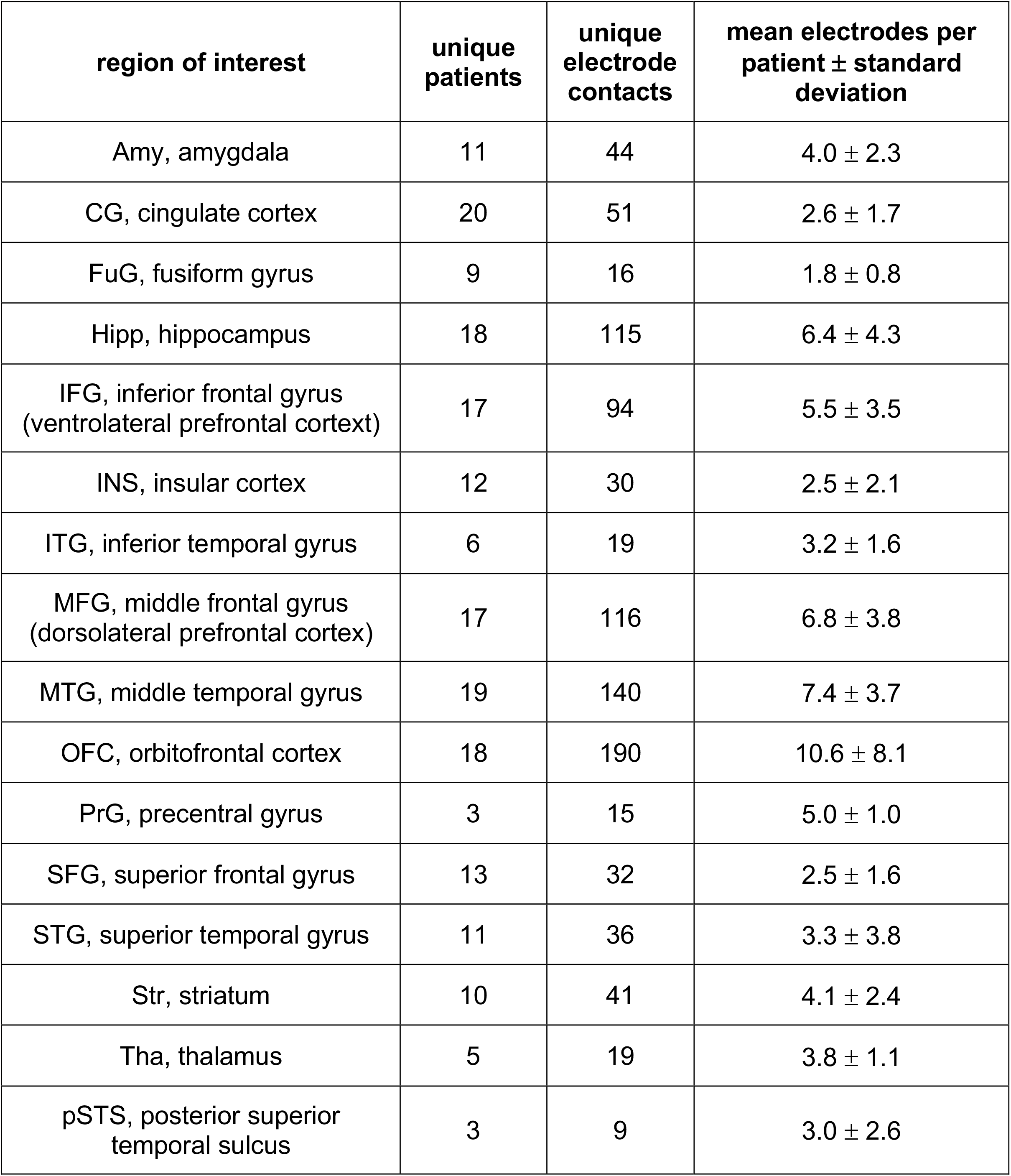
Brain regions sampled across twenty patients

| **region of interest** | **unique patients** | **unique electrode contacts** | **mean electrodes per patient ± standard deviation** |
| --- | --- | --- | --- |
| Amy, amygdala | 11 | 44 | $4.0 \pm 2.3$ |
| CG, cingulate cortex | 20 | 51 | $2.6 \pm 1.7$ |
| FuG, fusiform gyrus | 9 | 16 | $1.8 \pm 0.8$ |
| Hipp, hippocampus | 18 | 115 | $6.4 \pm 4.3$ |
| IFG, inferior frontal gyrus (ventrolateral prefrontal cortex) | 17 | 94 | $5.5 \pm 3.5$ |
| INS, insular cortex | 12 | 30 | $2.5 \pm 2.1$ |
| ITG, inferior temporal gyrus | 6 | 19 | $3.2 \pm 1.6$ |
| MFG, middle frontal gyrus (dorsolateral prefrontal cortex) | 17 | 116 | $6.8 \pm 3.8$ |
| MTG, middle temporal gyrus | 19 | 140 | $7.4 \pm 3.7$ |
| OFC, orbitofrontal cortex | 18 | 190 | $10.6 \pm 8.1$ |
| PrG, precentral gyrus | 3 | 15 | $5.0 \pm 1.0$ |
| SFG, superior frontal gyrus | 13 | 32 | $2.5 \pm 1.6$ |
| STG, superior temporal gyrus | 11 | 36 | $3.3 \pm 3.8$ |
| Str, striatum | 10 | 41 | $4.1 \pm 2.4$ |
| Tha, thalamus | 5 | 19 | $3.8 \pm 1.1$ |
| pSTS, posterior superior temporal sulcus | 3 | 9 | $3.0 \pm 2.6$ |

### Data collection and preprocessing

Neurophysiological data were recorded using a 128-channel data acquisition system (NeuroPort, Blackrock Microsystems, Salt Lake City, UT). Recorded data were bandpass filtered online at 0.3-250 Hz and sampled at 1 kHz. Reference channels were chosen for each patient consisting of a single intracranial depth electrode contact located in the white matter without artifact or epileptiform activity. The reference electrode was selected after electrode location labeling. An intracranial reference in white matter was chosen based on the improved signal quality over a scalp or skin reference.

A 60 Hz notch filter was applied to remove line noise. Recordings were referenced using the common median, which is more robust to large amplitude transient events than the common average reference (Rolston et al., 2009). Data were segmented into epochs of -250 ms to 1000 ms for the trial start and -1000 to 1000 ms at the trial end to facilitate analysis relative to specific trial events, which occurred at 0 ms. These events included: 1) initiation of stimulus delivery upon participant’s hand placement on the device and 2) removal of the participant’s hand from the device. After removing their hand from the device, participants responded via a touchscreen monitor to whether the stimulus was painful. Participants subsequently provided a numerical rating of perceived pain intensity on a scale of 0-10 (visual analog scale, VAS), which was not time-locked to the intracranial recordings. Both pain evaluation metrics were utilized as response variables in mixed-effects modeling.

Epileptiform discharges/noise spikes were identified as outliers if the epoch contained a value that was more than three median absolute deviations from the median. Channels were also visually inspected for artifacts after this automated analysis. Signals were downsampled to 500 Hz for subsequent analyses.

sEEG recordings allow evaluation of local neural activation through changes in amplitude within the broadband high-gamma frequency range (HFA; 70-150 Hz) in the recorded local field potential (LFP) (Ray and Maunsell, 2011; Buzsáki et al., 2012; Bartoli et al., 2019). HFA is highly correlated with both blood-oxygen-level-dependent fMRI and population firing rates. It is useful for bridging the gap between human neuroimaging results and findings from nonhuman primate electrophysiology studies (Miller, 2010). HFA was isolated through bandpass filtering the LFP between 70-150 Hz using a 4th-order zero-phase Butterworth filter. The absolute value of the Hilbert transform was applied to isolate the amplitude of the HFA component. The signal was smoothed using a moving average filter with a 300 ms fixed window. The epochs for each channel were normalized against baseline activity, using the mean and standard deviation from -3 to -2 s relative to the trial event onset. The time-locked epochs were z-scored using the mean and standard deviation to represent the relevant event being modeled.

### Experimental Design and Statistical Analyses

To investigate the relationship between HFA and painful experience, we used generalized linear mixed-effects (GLME) models. Trials from all patients were separated by brain region (see *Electrodes* for brain region labeling). We focused our analyses on the first and last seconds when the hand was exposed to the stimulus. We also evaluated the second after the hand was removed, right before they registered their binary choice. We chose a time window of 500 ms with a 250 ms overlapping sliding window to capture the likely physiological HFA of painful stimuli (Bastuji et al., 2016).

Two different types of GLME models were employed. The first model examined the relationship between HFA and whether the patient responded “yes” or “no” to whether the trial was painful (binary response). The fixed effect in this model was the mean HFA amplitude in one of the 16 brain regions during the 500 ms window, while the random effects were trial temperature and patient identity. A binomial distribution was used because the response variable was binary. The Wilkinson notation for this model was: y∼A+(A|B)+(A|C), where A was HFA amplitude for the particular brain region, B was the trial temperature on the thermoelectric device, and C was the patient identity. A random intercept and slope model was specified to account for variation in the fixed effect across subjects and temperature.

The second model evaluated the relationship between HFA and the numerical pain rating on a scale from 0 to 10. As in the first model, the fixed effect was the mean HFA amplitude in one of the 16 brain regions during a 500 ms window. As previously, the random effects were trial temperature and patient identity. This response variable was a numerical pain value between 0 and 10.

Statistical significance for each model was determined by a permutation test, where the model was re-evaluated 1,000 times with patients’ responses randomly permuted. The 95% range of the pseudo-t-statistics was used to determine the bounds of t-statistics associated with randomness. The true model was considered significant if the real t-statistic did not fall within these bounds. For the significantly predictive windows, an observed probability, *P*, was determined by adding the number of pseudo-t-statistics greater than the real t-statistic, multiplying by two to account for the two-sided distribution, and then dividing by the 1000 total observations. The smallest possible P-value was ≤0.002. Heatmaps of the real t-statistics were generated for each brain region and time window to visualize the models that significantly predicted the response variable.

### Evaluating model accuracy and predictions

We evaluated the accuracy of statistical models to predict the relationship between high-frequency activity (HFA) and pain response. The accuracy of the models was evaluated by comparing the real and predicted (fitted conditional) responses for each GLME. The error percentage was calculated as the absolute difference between the real and predicted responses that were most likely not painful, divided by the total number of trials and multiplied by 100. A lower error percentage indicates a more accurate model. To visualize the model’s accuracy, we made scatterplots of the fitted conditional VAS responses versus the real VAS responses for the brain regions.

Lastly, we evaluated the fixed-effects coefficients for all GLME models that predicted either the binary or VAS pain response for at least a one-time window. The coefficients were evaluated to understand how the response variable would change given a change in the mean z-scored HFA by one unit.

### Code Accessibility

Code used for preprocessing and analyses is available upon request.

## Results

We utilized intracranial recordings to analyze changes in HFA associated with thermal pain perception across a distributed brain network. GLME models examined the relationship between HFA changes and self-reported pain. Results showed significant changes in HFA amplitude in multiple brain regions, including the prefrontal cortex, lateral temporal cortex, and others, during the time windows surrounding the four trial events of the task.

### Relationship between broadband gamma and the binary pain response

We first evaluated the mean HFA amplitude during the first second the subjects’ hand was on the device (**Figure 2C**, “first second”). We found that the mean HFA amplitude in the ITG, superior frontal gyrus (SFG), and STG was significantly higher when subjects responded that the stimulus was painful (response = “yes”). An example of how the z-scored mean HFA for the “yes” responses was different from the “no” responses is shown in **Figure 2A**. In this instance, the mean HFA amplitude in the STG for the “yes” responses had a similar shape to the “no” responses. However, the “yes” maximum amplitude was greater than the “no” responses at the onset of the stimulus and the no response is delayed in time relative to the yes response.

**Figure 2:**
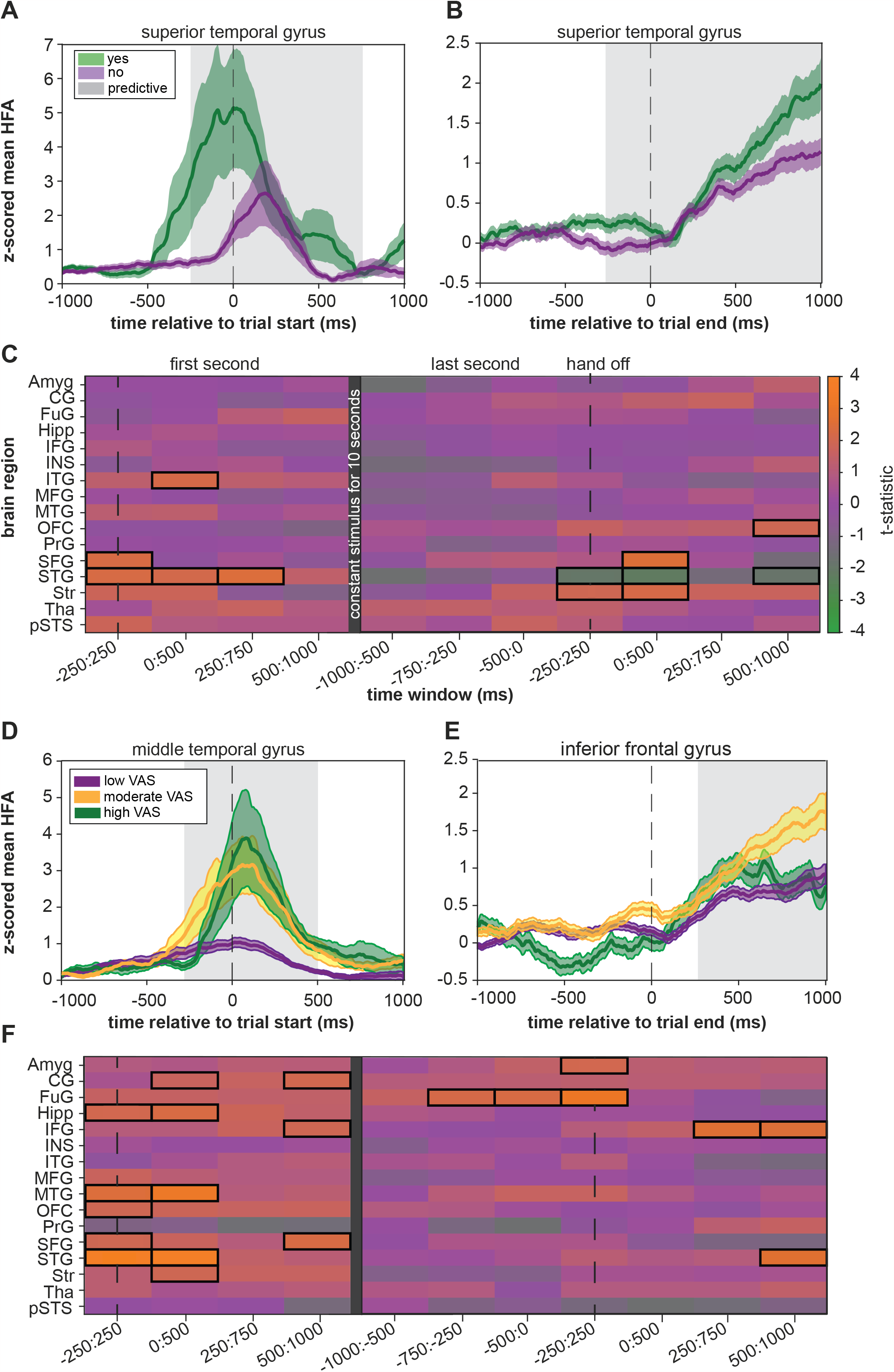
Relationship between HFA and the psychophysical evaluation of pain. 2A: The mean “yes” and “no” z-scored mean HFA during the first second of the trial within the superior temporal gyrus showed differential activity. The predictive time window from the heatmap is shaded in grey. 2B: At the end of the trial, there is still a time window where the HFA is predictive of the proportion of “yes” responses. However, the signal is different in shape and amplitude than at the onset of the stimulus. 2C: Heatmap coloring represents the real t-statistics for each brain region and overlapping sliding window for the binary response. Each row in the heatmap represents one of the sixteen brain regions analyzed. Each column represents one of the overlapping 500 ms windows that were analyzed. If the real t-statistic was significant (see *Statistical Analysis*), we outlined the region with a black box. The dotted vertical lines represent the onset and end of the trial. The SFG was predictive at the onset of the trial when the hand was initially on the device (−250:250 ms, P=0.006). The STG was also significant at the beginning of the stimulus until 750 ms (−250:-250 ms, P=0.006; 0-500 ms, P=0.02; 250-750 ms, P=0.004). The ITG was significant from 0:500 ms (P=0.034). When the hand was taken off, HFA increased in the OFC 500 to 1000 ms after hand removal from the device (P=0.034). The SFG (0:500 ms, P=0.004) and the striatum (−250:500 ms, P=0.028; 0:500 ms, P=0.016) also predicted this response but at different time windows. An increase in HFA in the STG was predictive of “no” responses (−250:250 ms, P=0.04; 0:500 ms, P=0.02; 500:1000 ms, P=0.05). 2D: The lower and upper third of the VAS ratings for all trials in the MTG were determined. The VAS values in the lower-third were classified as “low VAS.” The VAS ratings between one-third and two-thirds were classified as “moderate VAS,” and all ratings greater than the upper third were considered “high VAS.” The corresponding z-scored mean HFA for the three VAS categories demonstrates the HFA resulting in specific VAS responses. The high VAS scores have the largest HFA amplitude. 2E: Similar to Fig. 2D, but for the IFG. The mean HFA for all three groups seems relatively similar until 250 ms, where the HFA associated with the moderate VAS remains bigger, but the high VAS and low VAS values taper off. 2F: Heatmap coloring represents the GLME model results for determining whether HFA predicted the VAS response. Several unique regions in the first second of the stimulus included the cingulate gyrus (0:500 ms, P=0.044; 500:1000 ms, P=0.03), hippocampus (−250:250 ms, P=0.048; 0:500 ms, P=0.0240), IFG (500:1000 ms, P=0.032), MTG (−250:250 ms, P=0.006; 0:500 ms, P≤0.002), OFC (−250:250 ms, P=0.044), and striatum (0:500 ms, P=0.028). The SFG (−250:250 ms, P=0.02; 500:1000 ms, P=0.016) and the STG (−250:250 ms, P≤0.002; 0:500 ms,P≤0.002) were predictive in both models. Two areas demonstrated a relationship at the beginning and right after the stimulus— the IFG (250:750 ms, P=0.008; 500:100 ms, P=0.006) and the STG 500:1000 ms (P=0.022). The amygdala was involved in the time window of -250:250 ms (P=0.024), and the fusiform gyrus was involved in three-time windows (−750:-250 ms, P=0.024; - 500:0 ms, P=0.014; -250:250 ms, P≤0.002).

Subjects held their hand in place on the surface of the thermoelectric device for 10 seconds before removing it. We assessed which brain regions predicted the binary response at the end of the stimulus compared to the start by looking at the average HFA amplitude when subjects took their hand off the device (**Figure 2C**, “last second”). The results showed increased HFA in the orbitofrontal cortex (OFC) predicted “yes” responses after hand removal from the device. The SFG and the striatum also predicted this response at different time windows. An increase in HFA in the STG predicted “no” responses. While the HFA associated with “yes” responses was mostly higher in amplitude than “no” responses, it should be noted that visualizing the z-scored mean HFA across time, as in **Figure 2A/B**, does not account for interaction variables, such as temperature and patient identification even though the predictive regions did account for these variables.

We calculated an error metric for each brain region to evaluate the accuracy of each model’s prediction. The metric was the difference between the actual response and the response predicted by the model, with a lower error indicating a more accurate model. We only reported the accuracy for the predictive time windows from the analysis in **Figure 2C**. The model of the SFG was the most accurate at the start of the stimulus (**Figure 3A**). The STG had a slightly higher error. The STG was less accurate during the first 0 to 500 ms than the ITG. The STG was the least accurate of all models during the first second of the stimulus. The striatum was the least accurate when subjects removed their hand from the device but slightly less accurate than the STG. Over time, the accuracy of the STG improved, but the error of the striatum increased. The lowest error was found in the SFG, while the highest was in the striatum.

**Figure 3:**
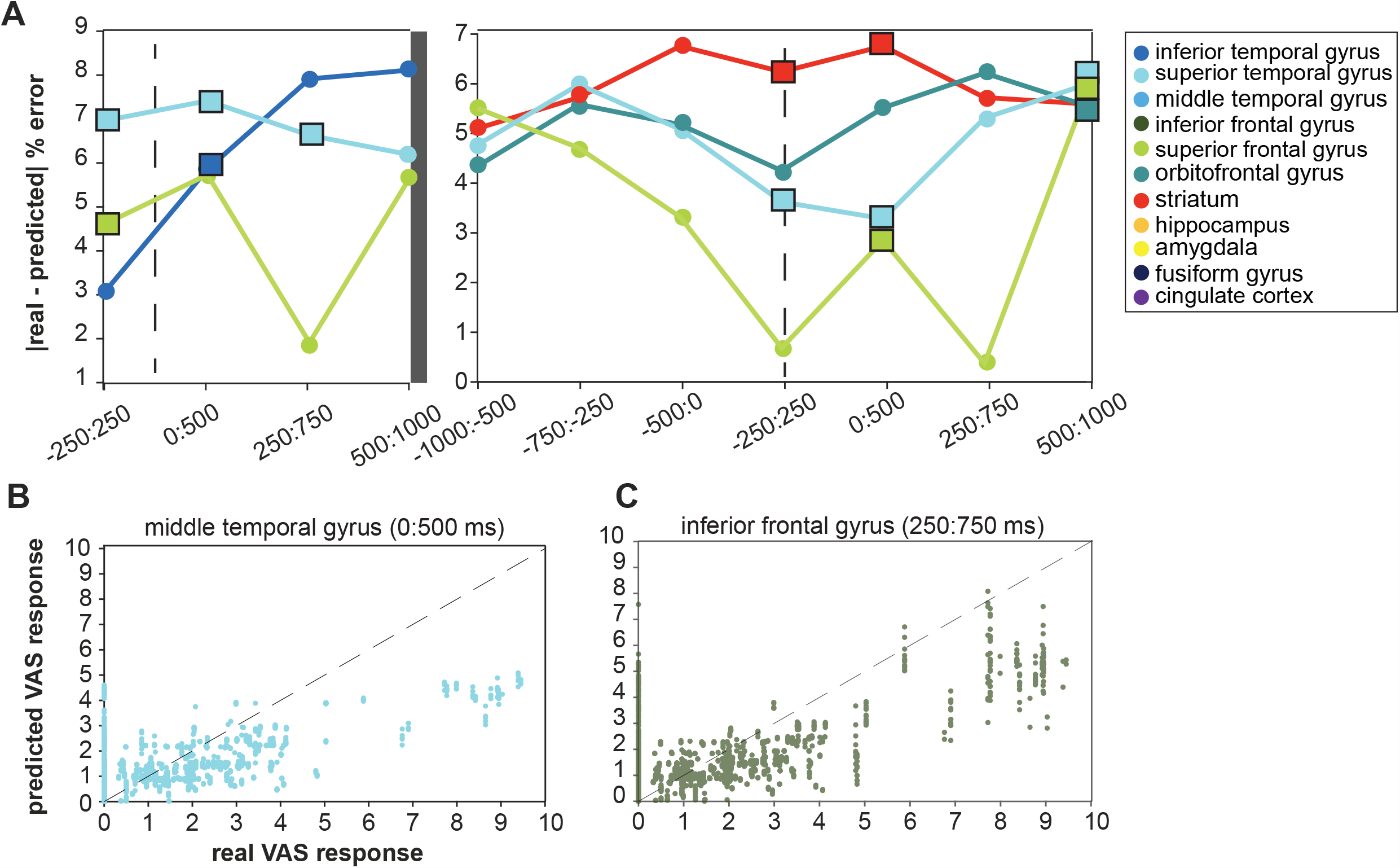
Assessing goodness-of-fit and fitted conditional responses of predictive GLME models. 3A: The absolute difference between the real and fitted conditional responses gave another metric for evaluating how well the model represented the binary responses. The SFG had the lowest error at the onset of the stimulus and the end of the stimulus. 3B: The fitted conditional responses versus the real VAS responses for the MTG during the 0:500 ms window show a proportional relationship. Not many real VAS responses were greater than five. 3C: The fitted conditional responses versus the real VAS responses for the IFG during the 250:750 ms window show a proportional relationship.

Lastly, we evaluated the relationship between the HFA and the binary response variables by looking at the estimated coefficient of the HFA models. This coefficient represented how much the response (“yes” or “no”) variable changed when the HFA changed by one unit (**Figure 4A**). The ITG had the largest coefficient in the first 500 ms of the stimulus, with a value of 0.12, which means that one unit increase in the HFA would result in a 12% increase in the proportion of “yes” responses. The SFG had the next biggest coefficient, and the superior temporal gyrus had the smallest coefficient.

**Figure 4:**
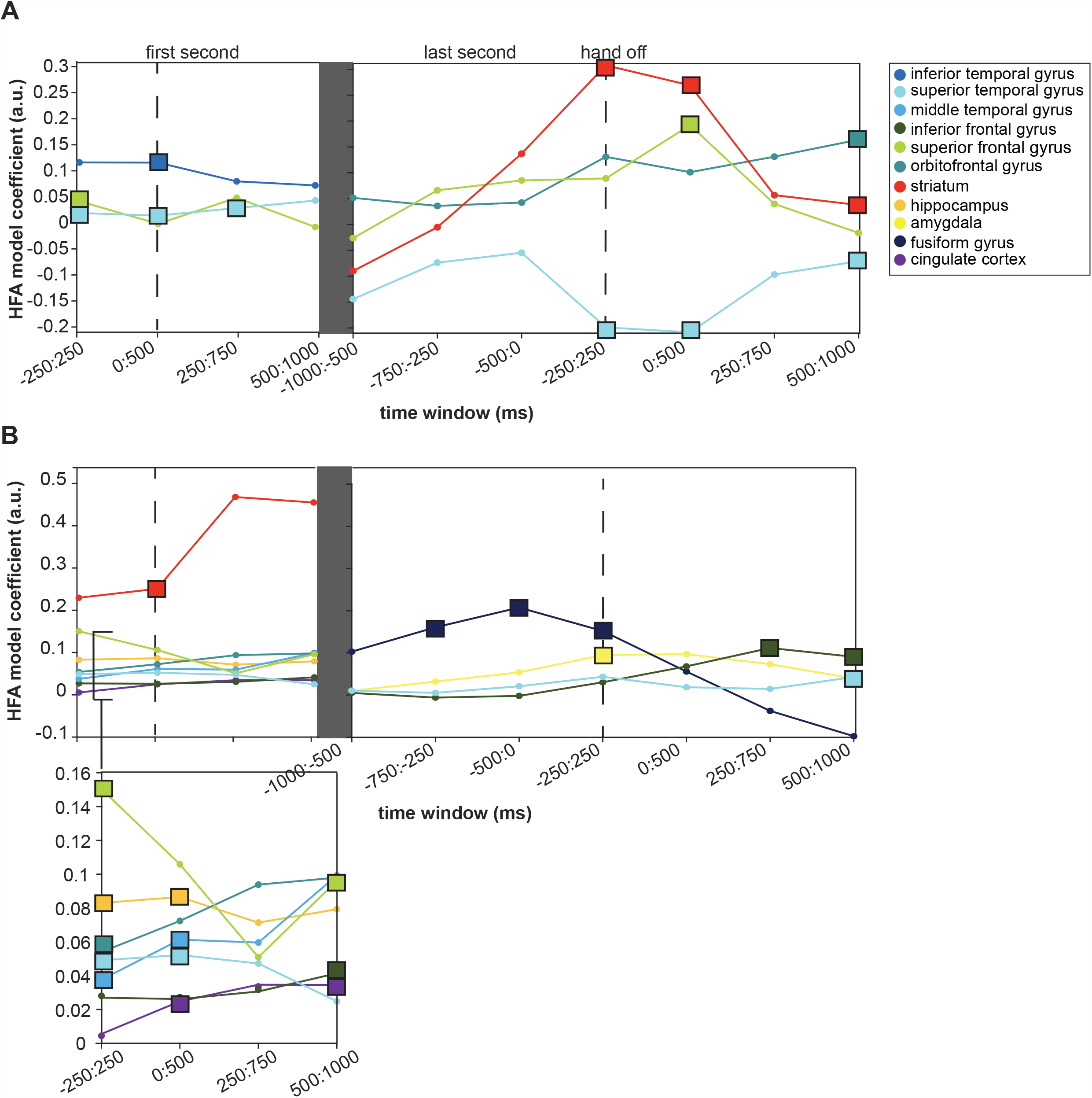
Model coefficient estimates predict change in psychophysical pain evaluation given HFA. 4A: The estimates for each time window and predictive brain region are shown. The square markers represent the time window that was predictive from the heatmaps in Fig. 2C. The STG had the smallest coefficient of 0.02, which means that a small increase in the HFA would lead to a 2% increase in the proportion of “yes” responses. The SFG had a slightly larger coefficient of 0.04 at the beginning of the trial (−250:250 ms). The coefficient for the STG increased slightly to 0.03. The largest coefficient was observed in the ITG from 0:500 ms (0.12). At the end of the stimulus, the SFG coefficient peaked right when the hand was taken off the device (0.19, 0:500 ms). The STG coefficient was negative throughout the trial, although slightly positive during the first second of the stimulus. The coefficient for the striatum was highest when the hand was removed from the device (0.31). It remained slightly higher than the coefficient for the SFG during this time. During the final time window, the OFC had the largest model coefficient at 0.16. The highest overall coefficient for the mixed-effect model with the binary response variable was from the striatum at -250:250 ms relative to the end of the stimulus (0.31). The second highest was from the SFG from 0:500 ms relative to the end of the stimulus (0.19). 4B: Model coefficients when VAS was the response variable were relatively similar between 0 and 0.1 during the onset. However, the striatum coefficient was much larger. During the first second of the stimulus, the biggest to smallest model coefficients were: striatum (0.25), SFG (0.15), hippocampus (0.087), MTG (0.061), OFC (0.054), STG (0.052), IFG (0.04), and the cingulate gyrus (0.035). The hippocampus, STG, MTG, and cingulate gyrus had increased coefficients in the first second. The coefficient for the SFG decreased over time. Although many regions had low coefficients (0.1), none were negative, which means increased HFA is linked to increased pain ratings. The fusiform gyrus had the largest coefficient of 0.21, with the peak occurring at the end of the stimulus. The IFG had a coefficient of 0.11, followed by the amygdala with 0.095 and the STG with 0.043. The peak model coefficient for the IFG was observed at 250 to 750 ms after the end of the stimulus.

In the last second of the stimulus, the model coefficient for the SFG remained stable. However, the coefficient peaked right when the hand was taken off the device. The STG coefficient was negative throughout the trial. In contrast, it was slightly positive during the first second of the stimulus. The coefficient for the striatum was highest at the end of the stimulus and remained slightly higher than the SFG coefficient. During the final time window, the OFC had the largest model coefficient. At the trial onset, the highest overall coefficient for the mixed-effect model with the binary response variable was from the striatum. The second highest was from the SFG from 0:500 ms relative to the end of the stimulus.

### Relationship between broadband gamma and the numerical rating of pain

We used mixed-effects models to study the relationship between the z-scored mean high-frequency activity (HFA) amplitude and the numerical pain ratings the subjects gave in response to the stimulus. The models were similar to the one used to evaluate the relationship between HFA and the binary pain response. However, in this case, the response variable was the numerical pain score participants assigned to each trial. During the first second of the stimulus, when each participant’s hand was on the device, we found that several brain areas had a significant relationship between HFA amplitude and the subjects’ pain rating **(Figure 2F)**. The t-statistic was positive for all the predictive regions, meaning that the z-scored mean HFA amplitudes were associated with increased pain ratings on the VAS.

Several regions uniquely predicted VAS response in the first second of the stimulus, including the cingulate gyrus, hippocampus, inferior frontal gyrus (IFG), MTG (**Figure 2D**), OFC, and striatum. The SFG and the STG were also predictive in the first second of the stimulus. Two cortical areas demonstrated a relationship between the mean HFA amplitude and pain ratings at the beginning and right after the stimulus— the IFG (**Figure 2E**) and the STG. However, many of the same brain regions were not involved when the stimulus ended. The amygdala was uniquely involved in the relationship between HFA and pain rating. Similarly, the fusiform gyrus was involved in three-time windows.

We tested the accuracy of our model by looking at the predicted and actual pain ratings on a scatter plot. As the pain stimulus was mild, most of the actual pain ratings were between 0 and 4, as shown in **Figures 3B and C**. When the real pain rating was around one, the predicted response was around one. However, when the actual pain rating was two, the predicted response seemed lower than two. This relationship suggests that the model may be under-predicting the pain score, but it could also be due to the limited number of higher real pain ratings.

To see how the HFA amplitude affects future predictions of pain, we looked at the model coefficients for each predictive region (**Figure 4B**). During the first second of the stimulus, the biggest to smallest model coefficients were: the striatum, SFG, hippocampus, MTG, OFC, STG, IFG, and the cingulate gyrus. The hippocampus, STG, MTG, and cingulate gyrus had increased coefficients in the first second. The coefficient for the SFG decreased over time. Although many regions had low coefficients, none were negative, which means increased HFA was always linked to increased VAS ratings.

We compared the models for the different brain regions after removing the hand from the device. The fusiform gyrus had the largest coefficient, with the peak occurring at the end of the stimulus. The IFG was the next largest, followed by the amygdala and the STG. The peak model coefficient for the IFG was observed at 250 to 750 ms after the end of the stimulus.

## Discussion

Mapping neural activity associated with ongoing pain with high spatial and temporal resolution is especially valuable and currently limited. Prior studies have used laser-based or thermode stimuli paired with noninvasive imaging or electroencephalography (EEG). Noninvasive imaging has provided valuable network-level information on whole-brain activity. However, it is limited in its temporal resolution. EEG can measure brain activity with a better temporal resolution but has limited spatial resolution of deep cerebral structures. Our approach involved acquiring recordings using intracranial stereo-EEG (sEEG) during a psychophysical pain testing paradigm to evaluate temporal changes in numerous regions involved in detecting, processing, and responding to painful stimuli (Caston et al., 2023). We used generalized mixed effects modeling to understand what regions underwent neuronal activations at the beginning of the stimulus, how those regions changed at the end, and how these neuronal activations were related to how the pain was reported.

Traditional models propose that a stimulus produces a percept and triggers a behavioral response (Romo, 2013). However, recent research indicates that this linear sequence may not apply to pain perception. May et al. argue that behavioral responses influence the perception of painful stimuli (May et al., 2017). This suggests that the cognitive and emotional aspects of pain processing are not separate from sensory perception but are integrated to create the subjective experience of pain. Schulz et al. have demonstrated that the subjective intensity of pain differs significantly from objective stimulus intensity and brief pain stimuli (Schulz et al., 2015). Chronic pain is a subjective experience that persists for extended periods. Therefore, in this study, we focused on the neural activity associated with the subjective experience of persistent pain.

Although there is limited research on temporal lobe abnormalities in chronic pain, it is believed that these areas are responsible for assigning emotional value to short-term memories associated with painful experiences (Godinho et al., 2006). Specifically, the STG plays a role in pain processing by monitoring the discrepancy between pain expectation and perception, anticipation, and expression (Budell et al., 2015; Palermo et al., 2015; de Pauw et al., 2019). In an fMRI study, Schwedt et al. (Schwedt et al., 2017) demonstrated that migraine patients may exhibit atypical connectivity between the MTG and a range of subcortical and cortical areas. Patients with chronic musculoskeletal pain disorders are similarly affected. For example, pain duration and intensity in patients with chronic musculoskeletal pain disorders correlate with decreased gray matter volume in the STG and MTG (Gerstner et al., 2011; Coppieters et al., 2016). Interestingly, recent evidence has shown that the STG plays a causal role in forming biased pain memories, as virtual lesions using transcranial magnetic stimulation reduced biased pain unpleasantness (Houde et al., 2020). The MTG and STG regions might contribute to the emotional response of a painful stimulus, leading to dysfunction in patients with chronic pain over time. To better understand where the pain changes are occurring within the MTG and STG, future analyses should use high resolution methods like sEEG. This is especially important since the STG is typically associated with speech and hearing, so a more precise approach can provide greater spatial specificity.

In our study, HFA at the onset of the stimulus in the ITG, SFG, and STG were significantly higher when participants responded that the stimulus was painful (“yes”). All three GLME models for these regions had a relatively low error. Since the ITG, STG, and MTG were predictive in both response types, we hypothesize that the temporal gyrus may represent more experiential aspects of pain.

At the end of the trial, increased STG HFA was associated with an increase in “no” responses, opposite to the trial start. We suggest that this change may reflect behavioral learning of the tonic stimulus or indicate a decrease in the neural processing demand, completion of the task, or the disengagement of attention from the task (Madhavan et al., 2015; Fu et al., 2018). Alternatively, it could also indicate inhibition or suppression of the brain region’s activity by other regions in the brain. All the coefficients for the predictive VAS models were positive, meaning that increased HFA was always linked to increased VAS ratings. An increase in HFA in more than one brain region near the same time suggests synchronized neural activity between these regions. The synchronized activity may indicate that these regions are involved in the same cognitive task or communicating. It would be valuable to study the connectivity of ongoing pain to understand the relationship between these neural features.

Other prefrontal cortical regions predicted both responses at the beginning and end of the stimuli. Prior groups have established the involvement of prefrontal cortices (e.g., IFG, MFG, SFG) in pain’s affective and cognitive dimensions (Salomons et al., 2007; Etkin et al., 2011; Kucyi and Davis, 2015; Bastuji et al., 2016). Recent studies also suggest that the medial prefrontal cortex encodes the subjective perception of ongoing pain and that gamma oscillations in this region encode subjection pain perception (Hashmi et al., 2013; Schulz et al., 2015). Time-frequency analyses in patients with chronic back pain revealed that ongoing pain intensity is reflected in prefrontal gamma oscillations (May et al., 2019). A recent study using EEG and a tonic pain stimulus in healthy patients observed a widespread increase of gamma power correlated with subjective pain intensity (Peng et al., 2014). Another study supported this finding, which found that gamma oscillations selectively encoded the subjective perception of tonic pain in the medial prefrontal cortex (Schulz et al., 2015). It is currently unclear whether the narrowband gamma activity occurs in the same regions or if the gamma responses are a true reflection of the neuronal populations in the medial PFC.

We show that the SFG and IFG encoded subjective perception at the beginning and end of the tonic stimulus. The prefrontal cortex receives ascending, nociceptive input and controls top-down pain (de Freitas et al., 2014; Ong et al., 2019; Kummer et al., 2020). Based on these regions exhibiting HFA at the start and end of the stimuli associated with the psychophysical correlates, we may have also seen this phenomenon. However, causal evidence of this mechanism in humans experiencing tonic pain is still needed.

More brain regions exhibited HFA activity parametrically related to the VAS response. At the beginning of the trial, HFA in the cingulate gyrus, hippocampus, orbitofrontal cortex, and striatum showed significant relationships with increased VAS ratings. By the end of the trial, HFA in the fusiform gyrus and amygdala were also associated with increased VAS ratings. While all these regions have previously shown involvement in pain processing, no one has shown a parametric relationship between behavioral responses and HFA during ongoing pain in these brain regions using intracranial electrodes. These findings suggest that the perception and psychophysical pain correlates depend on how patients are asked to evaluate it. The different brain regions involved in the VAS predictions highlight the importance of recognizing that brain dynamics can shift by changing just one aspect of the stimulus-perception-behavior relationship.

In this study, we utilized sEEG to study the spatiotemporal dynamics of tonic pain processing. While sEEG has great spatial and temporal benefits, several limitations exist. Electrode placement was determined based on the clinical hypothesis about the seizure onset zone, and there are innate differences in the sampled locations between patients. We combined electrode laterality and broadened specific electrode labels to obtain adequate statistical power for group-level analyses. Future analyses may use our findings to study the described regions with higher resolution. We also did our best to limit experimentation with patients receiving pain medication within 4 hours, despite pain medication being commonly indicated for patients with sEEG. Additionally, pain sensitivity may differ from one patient to another (at baseline and after surgical electrode placement), and sEEG is associated with lower pain medication use than other intracranial recording techniques (Scoville et al., 2021). To account for this, we used a maximum-likelihood adaptive procedure within the psychophysical testing paradigm to optimize the stimulus intensity for each patient. We also accounted for potential differences in stimulus intensity perception in our GLME models by including temperature as a random effect related to the HFA slope. Lastly, we observed the binary response model for the SFG and STG, and the VAS response model for the hippocampus, MTG, OFC, SFG, and STG were predictive starting at -250 ms relative to the stimulus onset. This could be explained by the small degree of error associated with the trial start recorded by the capacitive touch sensor. The patient may have their hand on the device but hasn’t applied enough pressure to trigger the recorded start time even though the stimulus is on.

Our study sheds light on the spatiotemporal dynamics of neural activity associated with psychophysical pain evaluation. Our findings indicate that depending on the psychometric variable in question, different brain regions may be involved in processing pain. These regions may exhibit distinct patterns of activity based on the reference frame (beginning or end of the stimulus). Additional research examining the temporal dynamics during the evolution of pain will enhance our comprehension of how the relationship between these regions changes as pain becomes a conscious perception.

## Acknowledgements

JDR was supported by the National Institute of Neurological Disorders and Stroke (K23 NS114178 and R21 NS113031). RMC was supported by the National Institute of Neurological Disorders and Stroke T32 NS115723. Thank you to Greg Stoddard for providing statistical consulting.

## Notes

### Competing Interest Statement

JDR has received consulting fees from Medtronic, NeuroPace, and Corlieve Therapeutics.

